# Opposite patterns of association of *TWIST1* expression with patient survival in SHH and Group 4 medulloblastoma

**DOI:** 10.1101/2025.03.19.644169

**Authors:** Karina Munhoz de Paula Alves Coelho, Matheus Dalmolin, Francis Rosseti Pedack, José Guilherme Pickler, Camila Barbosa, Julio Recalde, Rodrigo Blasius, Bruna Louise Silva, Marcelo A.C. Fernandes, Hercilio Fronza Junior, Paulo Henrique Condeixa de França, Rafael Roesler

## Abstract

**Purpose:** Twist1 is a transcription factor that regulates embryonic development, stemness, and differentiation, and can also stimulate initiation of tumorigenesis in peripheral solid cancers. However, its role in central nervous system tumors, including medulloblastoma (MB), the main type of malignant brain cancer that afflicts children, remains poorly understood. Here, we examined expression of Twist1, and its potential significance in prognosis, in different histological variants and molecular subgroups of MB.

**Methods:** Gene expression data for *TWIST1* and corresponding overall survival (OS) of patients was analyzed in 612 MB samples using a previously described dataset. A cross-sectional analysis of Twist1 protein content in 24 MB tumor samples from patients was carried out by immunohistochemistry.

**Results:** *TWIST1* transcript levels were higher in classic MB compared to desmoplastic tumors. Within samples with classic histology, higher *TWIST1* expression was associated with a longer OS. Tumors in the SHH subgroup had lower *TWIST1* expression compared to all other subgroups, and Group 4 showed lower expression than WNT tumors. In Group 4 MB, higher *TWIST1* levels were associated with shorter OS, whereas patients with SHH tumors and higher *TWIST1* levels showed longer OS. Twist1 protein was detectable in part of classic and LCA MB tumors belonging to the SHH, WNT or Group 3/4 subgroups.

**Conclusion:** We found opposite patterns of association between *TWST1* and patient survival in Group 4 and SHH MB subgroups. Our results highlight the importance of stratifying tumors by molecular subgroup and histological classification when exploring novel potential biomarkers and therapeutic targets in MB.

## Introduction

Medulloblastoma (MB) is the most common malignant brain tumor in children, posing a significant challenge due to its aggressive nature and long-term treatment consequences. MB arises in the cerebellum, probably from neural stem cells (NSCs) or granule neuron precursors (GNPs) which acquire a series of genetic and epigenetic alterations. While advances in classification have improved risk stratification, current treatment continues to rely on combined therapy comprising surgery, radiotherapy, and chemotherapy, approaches that, despite improving survival rates, often result in severe long-term side effects. A deeper understanding of the molecular drivers of each subgroup is essential for the development of more precise and less toxic therapies [1–4].

MB tumors can be classified on the basis of molecular or histological features. The four distinct molecular subgroups, namely wingless (WNT)-activated, sonic hedgehog (SHH)-activated, Group 3, and Group 4, differ in their underlying biology, patient outcomes, and therapeutic responses, with Group 3 being generally associated with the worst prognosis [5–9]. Histologically, MB can be separated according to morphology, nodularity, desmoplasia, and anaplasia into variants including classic, desmoplastic/nodular, and large cell/anaplastic (LCA). A special group of desmoplastic tumors is classified as MB with extensive nodularity (MBEN). Classic MBs are the most common, accounting for about 72% of tumors, and are characterized by relatively round nuclei, absence of increased cell size, and absence of frequent mitoses. Desmoplastic/nodular MB shows nodules of neuronal differentiation with intervening embryonal elements [10, 11]. Desmoplastic/nodular and MBEN MB have a better outcome in children whereas LCA tumors generally are more aggressive [12, 13].

Twist1 is a transcription factor that regulates differentiation of mesoderm tissues during embryonic development and can also act as an oncogene, promoting the formation of cancer stem cells and initiation of tumorigenesis [14–18]. In hematopoietic stem cells, Twist1 regulates the acquisition of stem cell-like properties and its expression declines as cells differentiate [19]. Twist1 interacts with β/δ-catenins to regulate fate transition in cranial neural crest cells [20], and, in the developing cerebellum, promotes the expression of microRNAs (miRNA) miR-199a and miR-214 to regulate the fate of specific neural cell populations [21]. In experimental models of peripheral solid tumor types including breast and prostate cancer, Twist1 is critical for stemness and metastasis, at least partially by promoting epithelial-mesenchymal transition (EMT) resulting in epigenetic reprogramming [22–24]

Given the role of Twist1 in regulating embryonic and cerebellar development, cancer stem cells, and tumorigenesis, in this study we aimed to examine Twist1 expression both at the mRNA and protein levels, and its potential significance in patient prognosis, in different histological variants and molecular subgroups of MB tumors.

## Methodology

### Gene expression

Gene expression data for *TWIST1* and corresponding survival information were obtained from the dataset described by Cavalli et al. [25] and are publicly available in the Gene Expression Omnibus (GEO) under the identifier GSE85217. Analysis was carried out as previously described [26, 27]. A total of 612 MB tumor samples with associated overall survival (OS) data were included in our analysis, histologically classified as follows: 337 samples with classic histology, 85 desmoplastic/nodular MB, 63 LCA MB, and 14 MBEN. In terms of molecular classification, the dataset comprised 113 samples from Group 3, 264 from Group 4 (G4), 172 from the SHH subgroup, and 64 from the WNT subgroup.

### Immunohistochemistry

We carried out a descriptive, cross-sectional analysis of MB tumor samples from patients from our local cohort. Histological variant and molecular subgroup were determined by immunohistochemistry. Twenty-four samples collected between January 2011 and February 2024 were included. Twist1 protein content was measured in tumor samples previously fixed in 10% formalin and stored in paraffin blocks. Samples in paraffin blocks with insufficient material for further testing were excluded. Antigenic recovery was performed using PT-Link equipment (Dako, Glostrup, Denmark) at high pH for 20 minutes at 95°C. Immunohistochemistry was performed using the Twist1 ‘clone 2C1a’ monoclonal antibody (Santa Cruz Biotechnology, Dallas, USA), diluted 1:50 in EnVision FLEX Antibody Diluent Solution (Dako) overnight. A 3% hydrogen peroxide solution (Dako) was used for 5 min for blocking peroxidase. EnVision FLEX Mouse LINKER solution (Dako) was added after incubation with the antibody and left for 15 minutes. Slides were incubated in EnVision FLEX/HRP polymer (Dako) for 20 min. EnVision FLEXDAB Chromogen (Dako) was used for 5 min for revelation. The slides were counterstained with Gill’s hematoxylin (Dako). Evaluation of nuclear expression of Twist1 was performed considering nuclear, cytoplasmic, and membranous staining. The slides were analyzed by two pathologists independently and blinded to clinical and histological data related to the samples. Molecular subgroup classification in these samples was determined by immunohistochemical detection of clinicopathological correlates GAB1, YAB1, and β-catenin [28].

### Statistical analysis

To investigate differences in gene expression, we applied the Wilcoxon test, followed by the Dunn test for pairwise comparisons. These analyses were performed for both histological variants and molecular subgroups. Statistical significance was determined using Holm-adjusted *p*-values, and all statistical tests were conducted using the ’ggstatsplot’ package. In addition, we assessed the correlation between gene expression levels and OS in MB patients. Patients were stratified into high- and low-expression groups using the ’Survminer’ package with the parameter ’minprop = 0.2’. Kaplan-Meier analyses of survival were performed using the ’Survival’ package [.

## Results

### *TWIST1* gene expression is higher and associated with better prognosis in MB with classic histology

*TWIST1* transcript levels were significantly higher in classic MB tumors compared to desmoplastic tumors (adjusted *p* value = 0.01, Fig. 1A). No significant differences in *TWIST1* expression were observed among other histological variants. Higher *TWIST1* expression was associated with a better prognosis indicated by longer OS in samples with classic histology (*p* = 0.039, Fig. 1B).

**Fig. 1.**
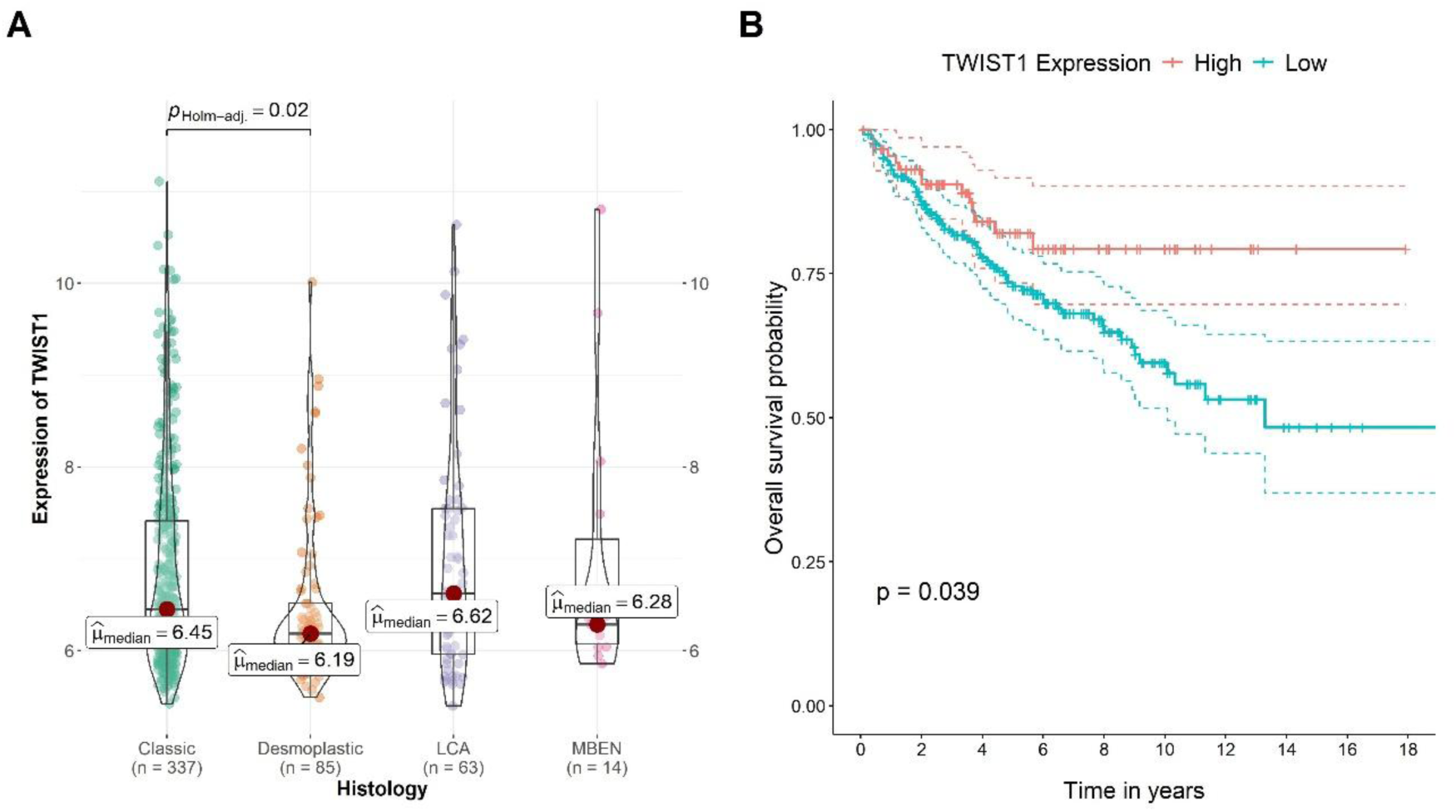
Gene expression of *TWIST1* in MB tumors classified according to histological variant. **A** *TWIST1* transcript levels in classic (*N* = 337), desmoplastic/nodular (*N* = 85), LCA (*N* = 337), and MBEN (*N* = 14) MB tumors from the Cavalli cohort [25]. **B** Kaplan-Meier analysis of OS in patients bearing MB tumors of the classic histology variant with low or high levels of *TWIST1* expression; *p* values are indicated in the figure.

### Group 4 and SHH MB show opposite patterns of association between *TWIST1* expression and prognosis

Analysis of *TWIST1* gene expression across the four molecular subgroups of MB tumors revealed that SHH tumors had lower *TWIST1* expression compared to all other subgroups. Group 4 MB exhibited significantly higher transcript levels in comparison with SHH tumors, but lower expression than the WNT subgroup. (Figure 2A). We further investigated the association of *TWIST1* expression with OS, and found that, in patients with Group 4 tumors, higher *TWIST1* levels were associated with worst prognosis indicated by shorter OS (*p* = 0.038, Fig. 2B), whereas in SHH 4 MB, patients with higher *TWIST1* expression had a more favorable prognosis as indicated by longer OS (*p* = 0.012, Fig. 2C).

**Fig. 2.**
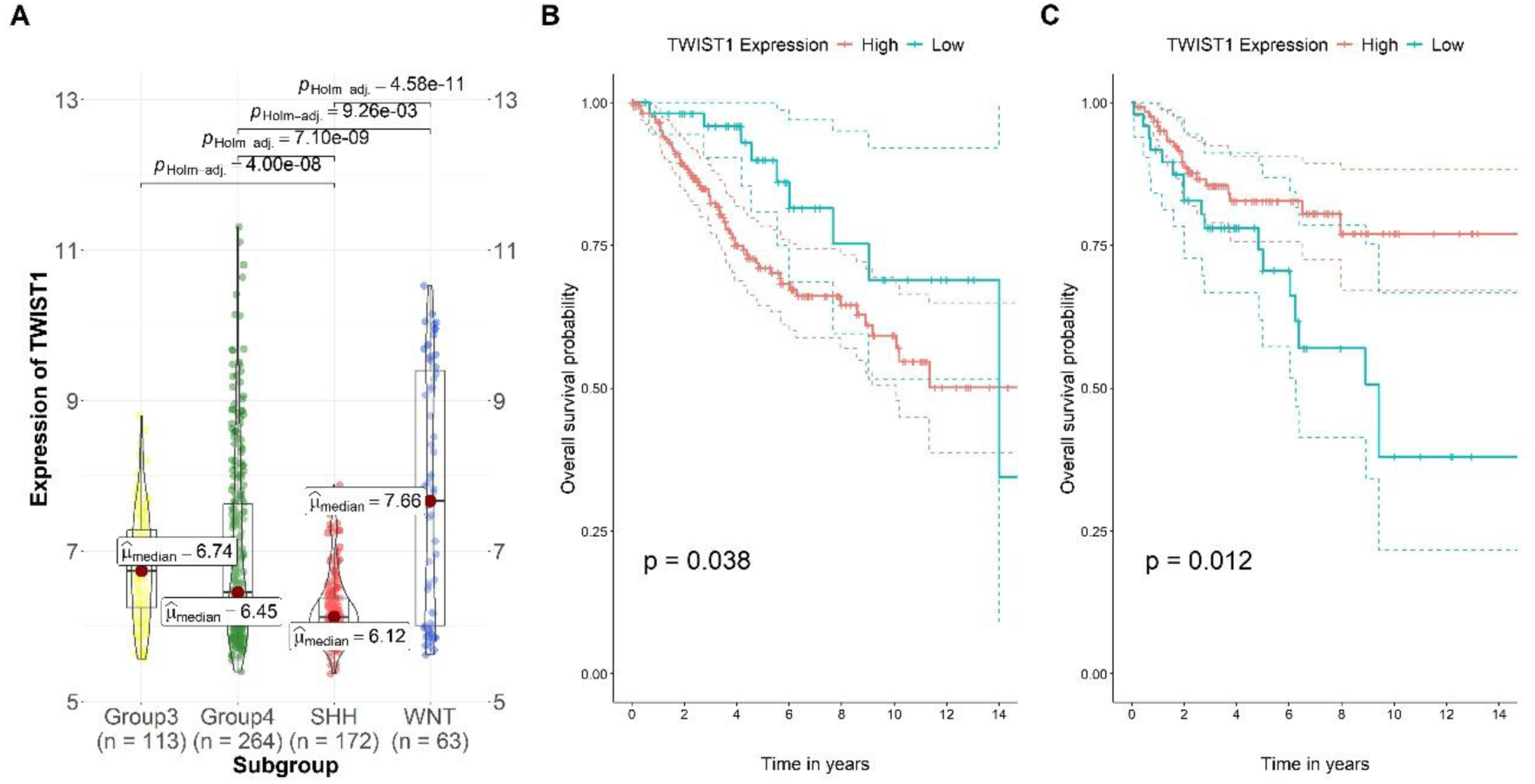
Gene expression of *TWIST1* in MB tumors classified according to molecular subgroup. **A** *TWIST1* transcript levels in Group 3 (*N* = 113), Group 4 (*N* = 264), SHH (*N* = 172), and WNT (*N* = 63) MB tumors from the Cavalli cohort [25]. **B** Kaplan-Meier analysis of OS in patients bearing Group 4 MB tumors with low or high levels of *TWIST1* expression. **C** Kaplan-Meier analysis of OS in patients bearing SHH MB tumors with low or high levels of *TWIST1* expression; *p* values are indicated in the figure.

### Twist1 protein is found in classic and LCA MB tumors belonging to the SHH, WNT, or Group 3/4 molecular subgroups

A total of 24 cases were analyzed, consisting of 12 females and 12 males, with ages ranging from 1 to 36 years. Fourteen (58.3%) cases were classified as Group 3 and 4, 6 (25%) as SHH, 1 (4,2%) as WNT, and 3 (12.5%) could not be classified, on the basis of clinicopathological correlates assessed by immunohistochemistry. In terms of histological variants, 15 (62.5%) cases were classified as classic, 2 (8.3%) as LCA, 6 (25%) as desmoplastic/nodular, and 1 (4.2%) as MBEN. Five (20.8%) cases were positive for Twist1 protein expression, whereas 19 (79.2%) were negative. These findings are detailed in Table 1. Representative images of MB tumors belonging to different histological variants and Twist1 immunohistochemical detection are shown in Fig. 3 and Fig. 4.

**Fig. 3.**
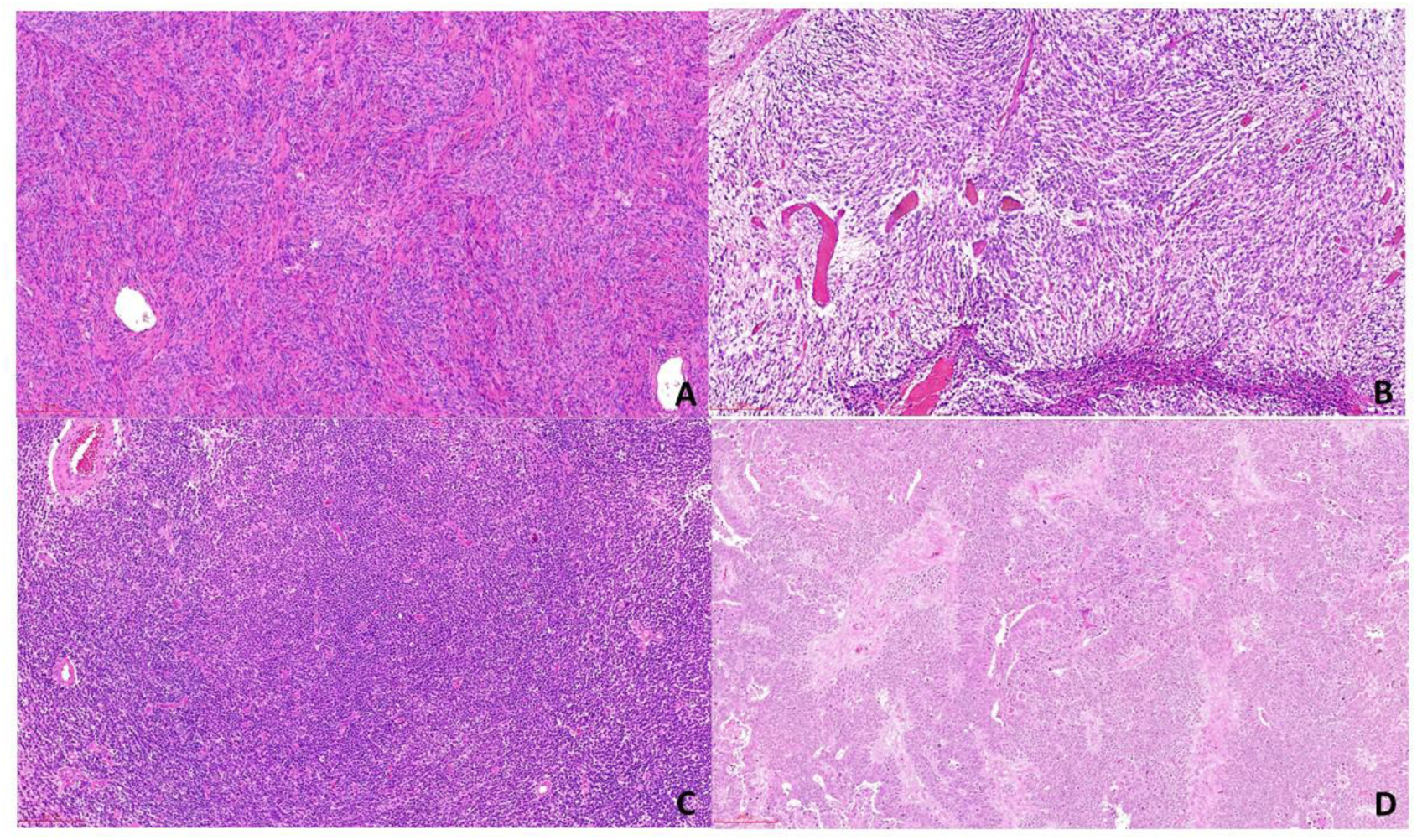
Representative histological images of MB tumors representing four histological variants. **A** desmoplastic/nodular, **B** MBEN, **C** classic histology, **D** LCA. Tumors were obtained from our local cohort. Hematoxylin and eosin (H&E) staining was used with 10 x magnification.

**Fig. 4.**
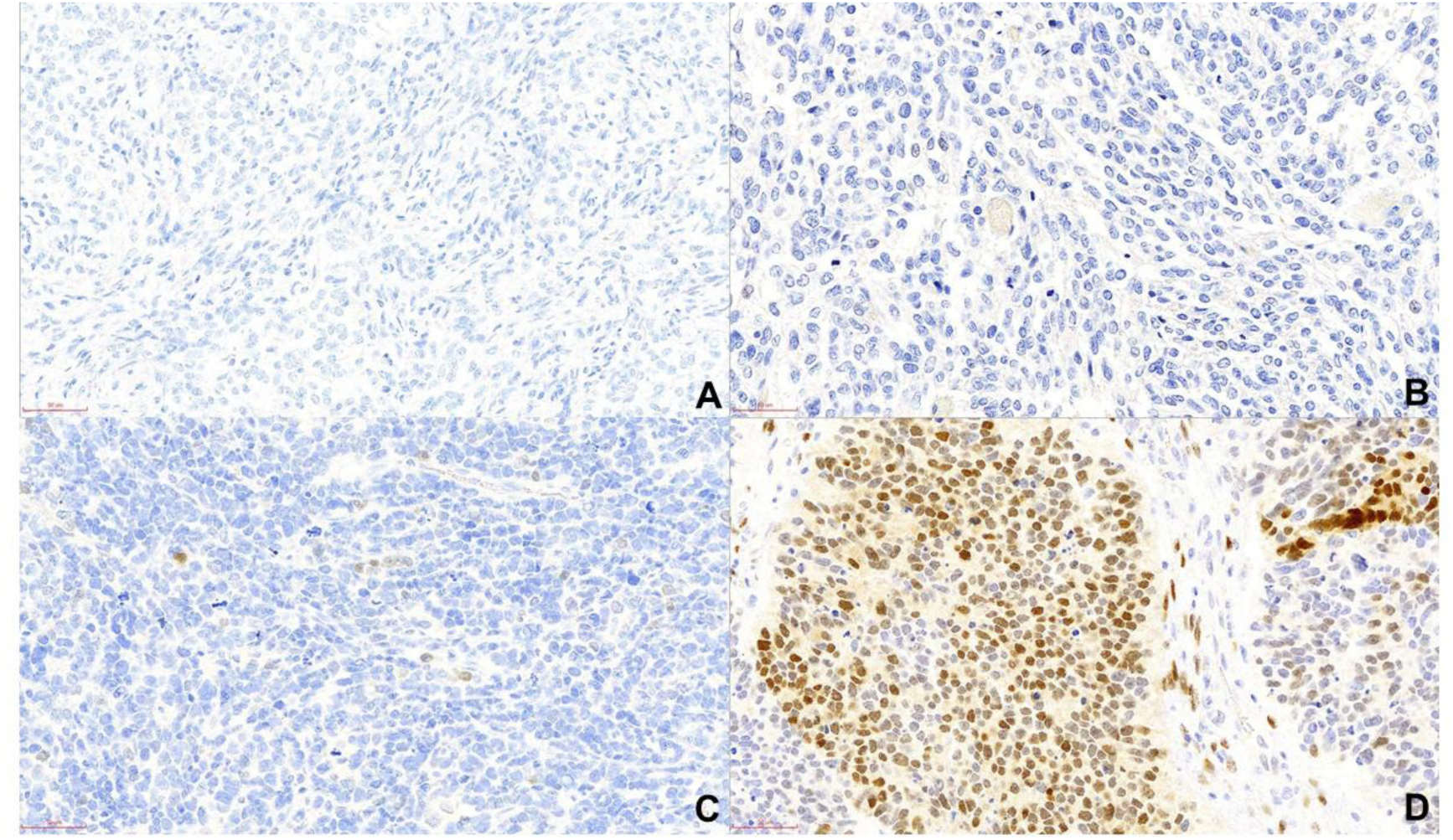
Representative histological images illustrating Twist1 protein expression or its absence **A** desmoplastic/nodular, Twist1-negative, **B** MBEN, Twist1-negative, **C** classic histology with weak and focal positivity, **D** LCA with strong and diffuse positivity. Tumors were obtained from our local cohort. Hematoxylin and eosin (H&E) staining was used with 10 x magnification.

**Table 1.**
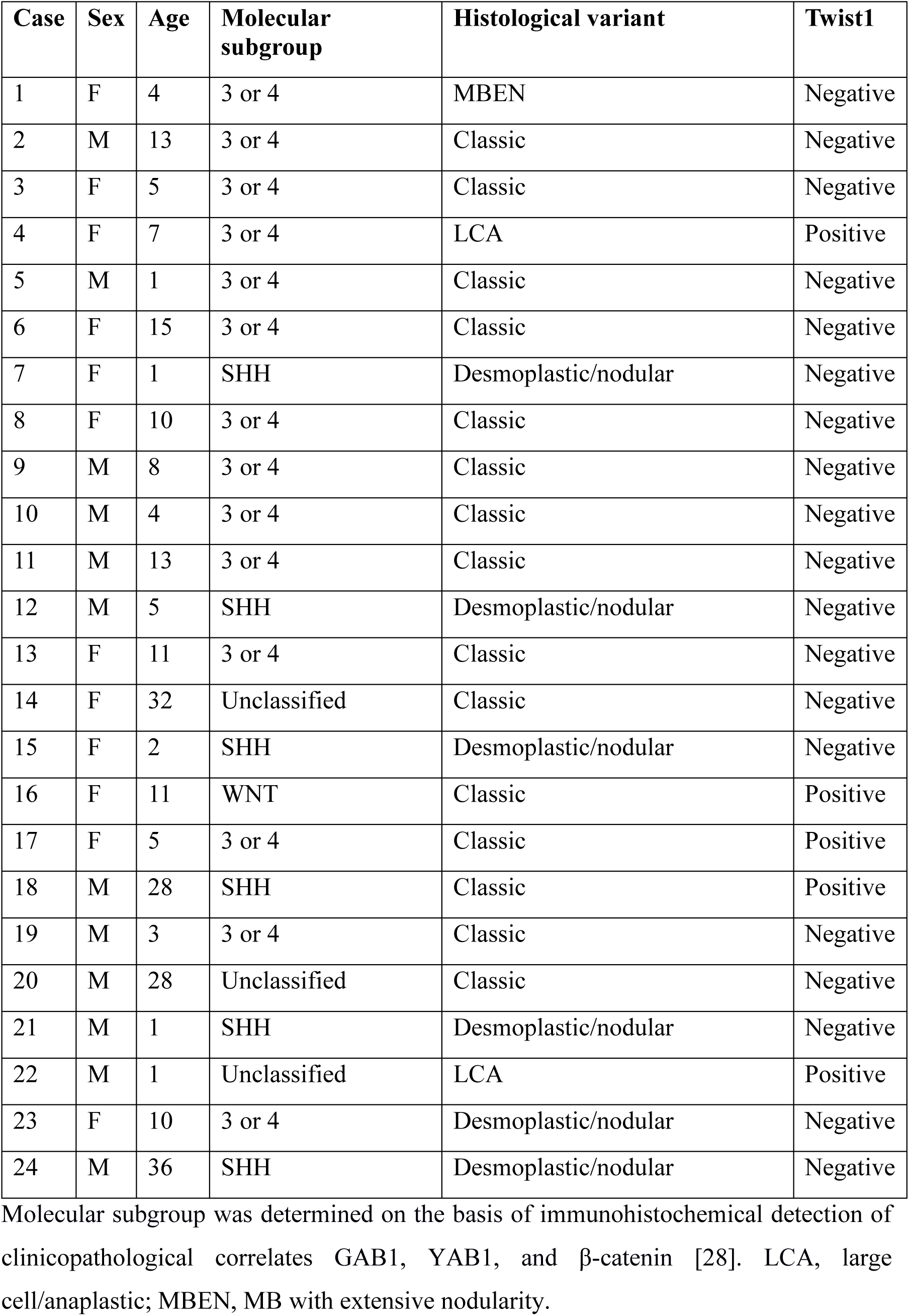
Immunohistochemical expression of Twist1 protein in MB tumors from a local cohort, classified into different molecular subgroups and histological variants.

| Case | Sex | Age | Molecular subgroup | Histological variant | Twist1 |
| --- | --- | --- | --- | --- | --- |
| 1 | F | 4 | 3 or 4 | MBEN | Negative |
| 2 | M | 13 | 3 or 4 | Classic | Negative |
| 3 | F | 5 | 3 or 4 | Classic | Negative |
| 4 | F | 7 | 3 or 4 | LCA | Positive |
| 5 | M | 1 | 3 or 4 | Classic | Negative |
| 6 | F | 15 | 3 or 4 | Classic | Negative |
| 7 | F | 1 | SHH | Desmoplastic/nodular | Negative |
| 8 | F | 10 | 3 or 4 | Classic | Negative |
| 9 | M | 8 | 3 or 4 | Classic | Negative |
| 10 | M | 4 | 3 or 4 | Classic | Negative |
| 11 | M | 13 | 3 or 4 | Classic | Negative |
| 12 | M | 5 | SHH | Desmoplastic/nodular | Negative |
| 13 | F | 11 | 3 or 4 | Classic | Negative |
| 14 | F | 32 | Unclassified | Classic | Negative |
| 15 | F | 2 | SHH | Desmoplastic/nodular | Negative |
| 16 | F | 11 | WNT | Classic | Positive |
| 17 | F | 5 | 3 or 4 | Classic | Positive |
| 18 | M | 28 | SHH | Classic | Positive |
| 19 | M | 3 | 3 or 4 | Classic | Negative |
| 20 | M | 28 | Unclassified | Classic | Negative |
| 21 | M | 1 | SHH | Desmoplastic/nodular | Negative |
| 22 | M | 1 | Unclassified | LCA | Positive |
| 23 | F | 10 | 3 or 4 | Desmoplastic/nodular | Negative |
| 24 | M | 36 | SHH | Desmoplastic/nodular | Negative |

Molecular subgroup was determined on the basis of immunohistochemical detection of clinicopathological correlates GAB1, YAB1, and β-catenin [28]. LCA, large cell/anaplastic; MBEN, MB with extensive nodularity.

## Discussion

Here, we report the first investigation of Twist1 expression and its association with prognosis in MB. We found that *TWIST1* transcript levels are particularly higher in MB tumors of the classic histological variant or belonging to the WNT molecular subgroup. In addition, *TWIST1* expression was associated in distinct ways with patient prognosis assessed by OS, depending on histological and molecular classification. Moreover, immunohistochemical analysis of Twist1 suggested that it can be found particularly in part of classic and LCA tumors belonging to the Group 3/4 or SHH subgroups, however the small sample number precludes a conclusive interpretation.

Only a few previous studies have addressed aspects of the possible Twist1 involvement in MB, which remains largely unknown. One report showed upregulation of Twist1 in a 3D Basement Membrane Extract (3D-BME) model of MB, and high nuclear Twist1 expression in the invasive edge of a MED1 orthotopic MB mouse model. Twist1 knockdown in MB cell lines resulted in reduced cell migration, whereas Twist1 binding to the promoter of the multidrug pump ABCB1 led to cell aggregation in metastatic and Twist1-overexpressing, non-metastatic MB cells. These findings suggest a role for Twist1 in promoting MB cell migration and possibly metastasis [29]. CD114-positive (CD114+) cells, which can be found in MB cell line cultures, patient-derived xenograft tumors, and primary patient tumor samples, and are possibly capable of evading chemotherapy, display slower growth rates and higher *TWIST1* expression levels compared to CD114- negative (CD114-) cells, raising the possibility that Twist1 affects resistance to treatment [30]. Neurogenic locus notch homolog protein 1 (Notch 1) signaling regulates both the initiation of metastasis and self-renewal of MB by stimulating the polycomb complex protein Bmi-1 through activation of Twist1 [31]. These findings provide early evidence that Twist1 influences aspects of MB tumorigenesis, progression, and metastasis, making it a target worth of further investigation.

Measuring mRNA expression of *TWIST1* in a series of 151 colorectal cancer samples showed that expression was restricted to tumor tissues and correlated with lymph node metastasis and shorter OS and disease-free survival in patients with stage I and stage II tumors [32]. Coexpression of Twist1 and the transcription factor ZEB2 measured by immunohistochemistry was associated with poorer OS in oral squamous cell carcinoma, particularly in patients without lymph node metastasis [33]. Immunohistochemical analysis of Twist1 also showed that overexpression is associated with histological grade, recurrence, and poor OS in patients with colorectal cancer [34, 35]. High Twist1 expression is also associated with shorter survival type I endometrial [36], pancreatic [37], and bladder [38], among other peripheral adult solid cancer types. In the pediatric cancer type Ewing sarcoma, Twist1 expression measured at either the mRNA or protein level was a negative prognostic marker for OS [39]. In neuroblastoma, the most common extracranial solid childhood cancer type, which arises from embryonal neural crest cells that later give rise to the sympathetic nervous system, Twist1 is a direct transcriptional target of the *MYCN* oncogene, which is associated with a worse prognosis [40]. Twist1 expression in NB patients is associated with poorer survival and metastasis. Moreover, CRISPR/Cas9 suppression of Twist1 in experimental NB reduces tumor growth and metastasis colonization in a mouse model [41]. Overall, these studies have also indicated that Twist1 is not necessarily ubiquitously expressed in normal and tumoral tissues – its protein expression can be low or even absent in most samples, consistently with what we observed in our immunohistochemical analysis.

Our finding in Group 4 MB tumors is consistent with the pattern of high Twist1 levels being predictive of poor prognosis reported in most previous cancer studies. However, we show that higher *TWIST1* expression is found in patients with longer OS bearing classic histology or SHH tumors. Within our results, this observation is particularly intriguing and encourages further studies aimed at exploring the role of Twist1 in classic MB in comparison with other histological variants, and Group4 versus SHH MB. As illustrated by a recent report showing that overexpression of ZIC1 suppresses the growth of Group 3 MB whereas in contrast it promotes the proliferation of SHH MB cells [42], transcription factors may play opposite roles in different molecular subgroups of MB.

## Conclusion

In summary, this report is the first to describe Twist1 expression at the mRNA and protein levels and associations of *TWIST1* gene expression with prognosis across different histological variants and molecular subgroups of MB. Remarkably, we found longer OS in patients with high-expressing classic MB, and opposite patterns of association between *TWIST1* levels and patient survival in the Group 4 and SHH tumor subgroups. Further studies should explore how Twist1 affects the initiation, progression, and metastasis in MB tumors belonging to distinct variants and subgroups.

## Author contribution

KMPAC, MD, FRP, JGP, PHCF, and RR designed the study. KMPAC, MD, FRP, CB, JR, RB, and BLS carried out analyses and analyzed the data. KMPAC, MACF, HFJ, PHCF, and RR provided materials and resources. KMPAC, MACF, HFJ, PHCF, and RR supervised the study. The first draft of the manuscript was written by KMPAC, MD, and RR. All authors revised previous versions of the manuscript. All authors read and approved the final manuscript.

## Data availability

The dataset analyzed in this study is available in the Gene Expression Omnibus repository, https://www.ncbi.nlm.nih.gov/geo/query/acc.cgi?acc=GSE85217.

## Declarations

## Ethics approval

This study was approved by the Institutional Research Ethics Committee as project number 6.839.450.

